# Time Lagged Multidimensional Pattern Connectivity (TL MDPC): An EEG/MEG Pattern Transformation Based Functional Connectivity Metric

**DOI:** 10.1101/2022.05.21.492913

**Authors:** Setareh Rahimi, Rebecca Jackson, Seyedeh-Rezvan Farahibozorg, Olaf Hauk

## Abstract

Functional and effective connectivity methods are essential to study the complex information flow in brain networks underlying human cognition. Only recently have connectivity methods begun to emerge that make use of the full multidimensional information contained in patterns of brain activation, rather than univariate summary measures of these patterns. To date, these methods have mostly been applied to fMRI data, and no method allows vertex-vertex transformation with the temporal specificity of EEG/MEG data. Here, we introduce time-lagged multidimensional pattern connectivity (TL-MDPC) as a novel bivariate functional connectivity metric for EEG/MEG research. TL-MDPC estimates the vertex-to-vertex transformations among multiple brain regions and across different latency ranges. It determines how well patterns in ROI *X* at time point *t*_*x*_ can linearly predict patterns of ROI *Y* at time point *t*_*y*_. In the present study, we use simulations to demonstrate TL-MDPC’s increased sensitivity to multidimensional effects compared to a univariate approach across realistic choices of number of trials and signal-to-noise ratio. We applied TL-MDPC, as well as its univariate counterpart, to an existing dataset varying the depth of semantic processing of visually presented words by contrasting a semantic decision and a lexical decision task. TL-MDPC detected significant effects beginning very early on, and showed stronger task modulations than the univariate approach, suggesting that it is capable of capturing more information. With TL-MDPC only, we observed rich connectivity between core semantic representation (left and right anterior temporal lobes) and semantic control (inferior frontal gyrus and posterior temporal cortex) areas with greater semantic demands. TL-MDPC is a promising approach to identify multidimensional connectivity patterns, typically missed by univariate approaches.

**Highlights:**

1. TL-MDPC is a multidimensional functional connectivity method for event-related EMEG
2. TL-MDPC captures both univariate and multidimensional connectivity
3. TL-MDPC yields both zero-lag and time-lagged dependencies
4. TL-MDPC produced richer connectivity than univariate approaches in a semantic task
5. TL-MDPC identified connectivity between the ATL hubs and semantic control regions

## 1 Introduction

Cognitive functions are generated by distributed networks of dynamically interacting brain regions (Bullmore and Sporns, 2009; Passingham et al., 2002). Understanding how cortical areas cooperate to produce complex cognition, requires assessment of their structural-anatomical connections and their functional interactions. A wealth of methods have been proposed to study connectivity from structural, functional and effective perspectives (Basti et al., 2020; Bastos and Schoffelen, 2016; Friston, 2011; Hipp et al., 2012; Lachaux et al., 1999; Le Bihan et al., 2001; Marinazzo et al., 2008; Nolte et al., 2008; Van Den Heuvel and Pol, 2010; Vinck et al., 2011). However, these methods are typically univariate, whilst functional interactions between brain regions are likely to be multidimensional (DiCarlo et al., 2012). Patterns of activity within brain regions may contain important information that is ignored in conventional univariate methods, in which dynamic patterns of brain activity are reduced to one value/time course by averaging or through principal component analysis (Anzellotti et al., 2017b, 2017a; Basti et al., 2019, 2018). Therefore, we need methods that can measure multidimensional (i.e. voxels-to-voxels) functional interactions between brain regions (Similar to Basti et al. (2020), we use the terms “multidimensional” to refer to scenarios where multiple time courses per brain region are explicitly taken into account. “Univariate” is used for cases where time courses within each region are collapsed across voxels (e.g. by taking the average), and “multivariate” and “bivariate” refer to cases of multiple or two univariate time courses for multiple or two brain regions, respectively.

Some of these issues have recently been addressed by the introduction of different multidimensional connectivity analyses (reviewed in Anzellotti and Coutanche, 2018; Basti et al., 2020). While multivariate activation methods exploit pattern information within regions-of-interest (ROIs) to assess information coding, such as Multi-Variate Pattern Analysis (MVPA) (Cichy and Pantazis, 2017; Haxby, 2012) or Representational Similarity Analysis (Karimi-Rouzbahani et al., 2022; Kriegeskorte et al., 2008; Laakso and Cottrell, 2000), multidimensional connectivity methods make use of the pattern-to-pattern relationships among different ROIs to identify functional interactions between brain areas. For example, Multivariate Pattern Dependence (MVPD) (Anzellotti 2017a) determines statistical dependencies between patterns of responses in different brain regions whose dimensionalities are first reduced by PCA. This is accomplished by testing for either linear or nonlinear relationships between principal components in different regions. Unlike univariate approaches, when more than one principal component is used, some pattern information is retained which is a step forward from univariate techniques. However, the use of PCA for dimensionality reduction in this approach obscures the original voxel-to-voxel relationships between activity patterns in different ROIs. This issue was addressed in fMRI data by Basti et al. (2019). These authors estimated the linear transformation matrices between patterns using cross-validated ridge regression. Assessing the goodness-of-fit, sparsity and pattern deformation allowed more detailed examination of these voxel-to-voxel transformation matrices.

The above-mentioned multidimensional connectivity methods have mostly focused on functional Magnetic Resonance Imaging (fMRI) which is well-known to have limited temporal resolution. Electroencephalography and magnetoencephalography (EEG and MEG) signals track processes at a timescale appropriate for perceptual and cognitive processes and, in combination with appropriate source estimation procedures, have reasonable spatial resolution for cortical activity. Thus, appropriate analysis methods would enable multidimensional connectivity among brain regions to be tracked across different stages of perceptual and cognitive processes. A few methods that exploit vertex-to-vertex relationships in EEG/MEG data have already been proposed, such as the Multivariate Interaction Measure (MIM, Ewald et al., 2012), multivariate lagged coherence (MVLagCoh, Pascual-Marqui, 2007a), and multivariate phase-slope-index (MPSI, Basti et al., 2018). These are frequency-domain methods that expand the imaginary part of coherency (ImCoh, Nolte et al., 2004), lagged coherence (Pascual-Marqui, 2007b), and phase-slope index (Nolte et al., 2008), respectively, to multivariate time-series. Frequency domain methods depend on the choice of valid frequency bands as well as latency ranges to estimate the spectral connectivity metrics (Bastos and Schoffelen, 2016). While there is ample evidence that some brain processes and brain states are reflected in specific frequency bands (Fries, 2015; Siegel et al., 2012), it is uncertain that this generally the case, as for example for short-lived brain processes in event-related experimental paradigms. To date there is no multidimensional connectivity method utilizing EEG/MEG data to estimate the vertex-to-vertex transformations of activity patterns across time. Here, we fill this gap by extending the approach of Basti et al. (2019) to EEG/MEG source estimates.

More specifically, we propose Time-lagged Multidimensional Pattern Connectivity (TL-MDPC) as a novel functional connectivity method for event-related data that not only estimates relationships between the spatial activity patterns of two ROIs (as in Anzellotti et al., 2017a; Basti et al., 2019), but also examines their relationship over time through estimating vertex-to-vertex transformations for pairs of time points.. The aim of this method is to determine how well patterns of ROI *X* at time point *t*_*x*_ can predict patterns of ROI *Y* at time point *t*_*y*_ through a linear transformation. The cross-validated goodness of fit of this transformation acts as a connectivity metric for every pair of ROIs and time points. This results in a bivariate undirected functional connectivity metric, and thus shares the limitations of other exemplars of this category, such as coherence or phase-locking values (Bastos and Schoffelen, 2016). It can detect statistical relationships between signals across trials in event-related experimental designs, but does not allow inferences about direction and causality of the underlying effects. It does not require a separation of signals into frequency bands and is relatively computationally inexpensive. It also furnishes a matrix relating the response pattern in ROI *X* at each time point to ROI *Y* at each time point, which can be interrogated to discover the nature of the transformation between patterns in different regions over time (Basti et al., 2019), even if the patterns themselves change dynamically.

Compared to activity patterns in fMRI data, EEG/MEG source estimates have a limited spatial resolution and are inherently smooth (Hauk et al., 2019; Molins et al., 2008; Samuelsson et al., 2021). Therefore, ROI patterns in source estimates contain redundant information. We used a novel approach to determine the most informative vertices in ROIs using k-means clustering of patterns across trials and selecting the vertex with the highest variance in each cluster. Reducing ROIs to their most informative vertices significantly increases computational processing speed. However, unlike a PCA approach, this approach does not conflate activity across voxels, and it does not require a pre-definition of latency ranges, i.e., it can be applied on a sample-by-sample basis. Thus, our approach allows the estimation of pattern transformations between specific vertices in two regions and for specific pairs of latencies. Following this reduction of patterns, the multidimensional relationship between patterns is estimated using cross-validated Ridge Regression (Hoerl and Kennard, 1970), i.e., potential overfitting is avoided using 10-fold cross validation and regularization of the underdetermined regression problem. The explained variance (EV) from the transformation between patterns is used as the goodness-of-fit. High EV indicates a strong linear relationship between the activity patterns of two ROIs at two latencies.

We evaluate our novel approach in simulations, as well as in real EEG/MEG data. Our simulations were designed to demonstrate that 1) TL-MDPC is indeed sensitive to linear multidimensional relationships between patterns, while a corresponding univariate version of this method is not; 2) TL-MDPC is also sensitive to univariate relationships, and 3) TL-MDPC is not prone to producing false positives for independent random patterns. We demonstrate these findings in simulations with typical pattern dimensions and numbers of trials, as well as over a wide range of SNRs.

In our EEG/MEG data analysis, we address timely questions about semantic brain networks, following our recent study using evoked responses and coherence analysis (Rahimi et al., 2022). Previous literature on semantic networks has linked the anterior temporal lobe (ATL) to semantic representation (Acosta-Cabronero et al., 2011; Binder et al., 2016; Martin, 2016; Pobric et al., 2007; Rogers et al., 2004) and posterior temporal cortex (PTC) and inferior frontal gyrus (IFG) to semantic control (Badre et al., 2005; Jackson, 2021; Jefferies and Lambon Ralph, 2006; Lambon Ralph et al., 2016; Noonan et al., 2013). The role of angular gyrus (AG), another region often associated with semantics, is less clear as it has been implicated in semantic representation (Binder et al., 2009), control (Noonan et al., 2013) and episodic memory processes (Humphreys et al., 2015). While the semantic network is usually reported to be left-lateralised (Binder et al., 2009), the ATL displays a graded lateralisation across both cerebral hemispheres depending on stimulus and task features (Marinkovic et al., 2003; Olson et al., 2007; Patterson et al., 2007; Pobric et al., 2007; Rice et al., 2015b, 2015a; Visser et al., 2010). Therefore, our ROIs comprise left-hemispheric regions commonly assumed to be involved in word recognition and semantics (left ATL, PTC, IFG, AG, primary visual area (PVA)), as well as right ATL. In general, evidence on semantic networks from dynamic neuroimaging methods is still scarce (Rahimi et al., 2022). Our previous study (Rahimi et al., 2022) compared tasks with varying semantic demands, revealing a critical role for left ATL in the semantic network. Utilising spectral coherence as a conventional univariate connectivity method, we found rich connectivity for left ATL, identifying connections with frontal semantic control areas as well as right ATL. However, the absence of connectivity between other nodes of the semantic network could be due to the fact the coherence is only sensitive to a very specific type of statistical relationship in specific frequency ranges. Utilising the same dataset, here we probe for further functional connectivity in the semantic network by comparing TL-MDPC with its univariate counterpart, time-lagged univariate connectivity (TL-UVC). The connectivity score for each pair of ROIs at each latency is presented in a time-time matrix which we call a temporal transformation matrix (TTM). This comparison demonstrates the utility of the TL-MDPC method in real brain data and confirms and extends the results of our previous study by elucidating the dynamic connectivity of the semantic network.

## 2 Materials and Methods

### 2.1 Time-Lagged Multidimensional Pattern Connectivity (TL-MDPC)

#### 2.1.1 General idea

The relationship between multidimensional time series is often investigated using multivariate autoregressive models (Anderson et al., 1998; Ding et al., 2000; Dutta et al., 2018; Schlögl, 2000; Schlögl and Supp, 2006; Siggiridou and Kugiumtzis, 2015; Zhang et al., 2017). The number of coefficients in the transformation matrix between two multidimensional time series is larger than the number of time series (in our case the numbers of vertices in an ROI). Thus, the estimation of these matrices from one or even a few samples is underdetermined. For continuous time series, this is addressed by estimating the coefficients across all samples in a time series (or a large enough subset), assuming that the matrix is stable across the whole time series. This may, for example, be suitable for resting state data (Blinowska et al., 2017; Colclough et al., 2015; Liegeois et al., 2017; Olejarczyk et al., 2017). However, event-related data is unlikely to be stable, since surface topographies and source distributions can change within tens of milliseconds. Nevertheless, the assumption of event-related designs is that brain processes are similar across a number of trials, at least for stimuli of the same category and in the same task. Thus, the transformation matrices can be estimated for pairs of ROIs and pairs of latencies across trials. The resulting problem may still be underdetermined and ill-posed, and may therefore require a regularization procedure. This approach was applied to fMRI data by Basti et al. (2019). Here we extend and apply it to EEG/MEG data as TL-MDPC as follows.

In order to deal with event-related data, let us consider *X* and *Y* as matrices with activity patterns in ROI X and ROI Y at time point *t*_*x*_and *t*_*y*_ of size *n*_*t*_ × *n*_*X*_ and *n*_*t*_ × *n*_*Y*_, respectively, where *n*_*t*_ is the number of trials, and *n*_*X*_ and *n*_*Y*_ are the number of vertices in the two ROIs. Figure 1 illustrates patterns of activity in ROI X and ROI Y over time. We are interested in whether there is an all-to-all mapping between the vertices (or voxels) of the patterns of the two ROIs at each pairs of time points. In other words, we want to see how well patterns of ROI X at time point *t*_*x*_ can predict patterns of ROI Y at time point *t*_*y*_ through a linear transformation. We will use the explained variance of this transformation as a connectivity metric. In order to avoid overfitting, the explained variance will be obtained via cross-validated regularized ridge regression.

**Figure 1.**
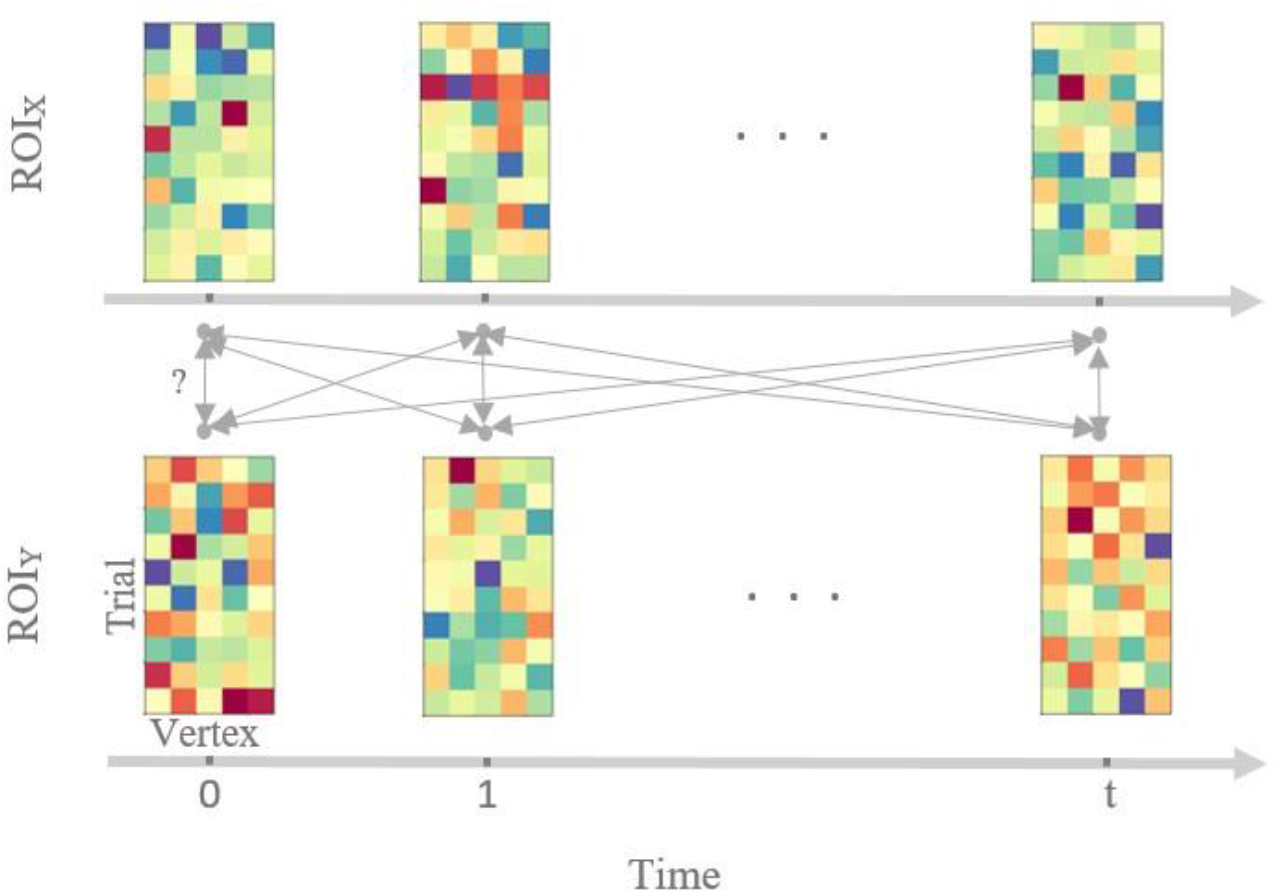
Illustration of TL-MDPC for event-related activity patterns in two ROIs, ROI X and ROI Y, across trials and across latencies. Each matrix represents activity patterns in one ROI at one latency, with rows representing activity patterns across different trials, and columns representing different vertices in the ROI. We test whether there is an all-to-all mapping between the patterns of the two ROIs at different latencies and determine how well *X* and *Y* can predict each other, by estimating a linear transformation per ROI pair and latency pair. Bidirectional arrows indicate possible transformations between pairs of patterns.

### 2.1.2 TL-MDPC: Modelling statistical dependence

We can estimate the transformation *T* from *X* to *Y* through Ridge Regression (Hoerl and Kennard, 1970) using a train subset of trials:

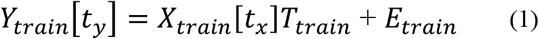

where *T*_*train*_ is the transformation matrix, of size *n*_*x*_× *n*_*y*_ and *E*_*train*_ is the error matrix of size *n*_*t*_ × *n*_*y*_. *T*_*train*_ can be estimated using the regularized pseudoinverse of *X*_*train*_:

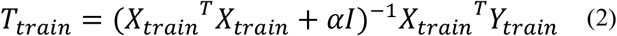

Where *α* is the regularization parameter (Tikhonov and Arsenin, 1977) to be determined using cross-validation and *I* is the Identity matrix of size *n*_*x*_× *n*_*x*_.

After estimating the coefficient matrix *T*_*train*_, ROI *Y*’s predicted patterns can be obtained using the test subset and *T*_*train*_ as follows:

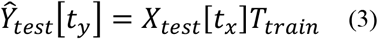

where *Ŷ*_*test*_ is the predicted pattern. For each vertex j=*1*,…, *n*_*y*_, we then compute the explained variance (EV):

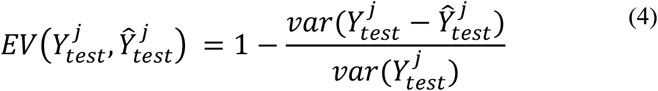

Finally, as a metric for multidimensional connectivity between two ROIs, we average the EV across vertices:

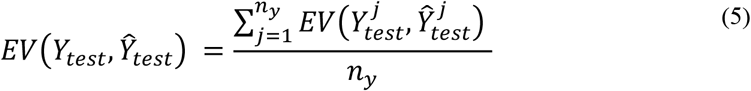

Thus, the best possible score in case of a perfect linear relationship between patterns is 1, whereas the score would approach zero or can even be negative when there is no linear relationship. This measure of explained variance can produce negative values if the magnitude of the prediction error is larger than the magnitude of the data themselves. This indicates that the model predicts incorrect data. In our own simulations and real data, we indeed observed some small negative values, which we do not consider interpretable. Thus, we replaced negative values with zeros, i.e., we reported *max*(*EV*(*Y*_*test*_, Ŷ_*test*_), 0) as the final metric.

TL-MDPC can be computed in two ways for each ROIs pair: *Y* can be predicted from *X* and *X* can be predicted from *Y*. The transformations in the two cases are unlikely to be the same. Indeed, this is impossible if the numbers of vertices in the two regions are different. Because our metric is not directional and only reflects a statistical relationship between patterns, and to avoid any potential bias due to different number of vertices and noise, we compute transformations for both cases and use the average EV as the connectivity metric.

### 2.1.3 Cluster-based spatial sub-sampling of vertices within ROIs

Given the limited spatial resolution of EEG/MEG signals, not all vertices within an ROI are independent (they “leak” into each other) (Hauk et al., 2019; Palva et al., 2018). In other words, activity patterns across vertices within ROIs are smooth, and including all vertices in the above transformation estimation procedure might lead to a large amount of redundant information. Transformations between this redundant information would be computationally expensive with little gain. Here, we propose a novel preprocessing method based on unsupervised k-means clustering to spatially sub-sample vertices within an ROI in order to determine the optimal amount of informative vertices needed to describe the activity pattern in an ROI.

The aim is to partition all vertices into k clusters, such that each cluster contains vertices with similar activation profiles across trials. To find the optimum number of clusters, we use the *elbow method* (Ng, 2012) implemented in Python^1^. After clustering, we pick the vertex with the highest variance from each cluster as the cluster representative. This removes redundant vertices and reduces the dimension of patterns and subsequent vertex-to-vertex transformation. Our patterns in ROI *X* and Y at time points *t*_*x*_and *t*_*y*_ are now of reduced size *n*_*t*_ × *n*_*x*_and *n*_*t*_ × *n*_*y*_, where *n*_*x*_and *n*_*y*_ are the number of clusters in *X* and *Y*. Prior to this step, we standardised each pattern matrix per ROI and time point by demeaning and variance normalization.

#### 2.1.4 Univariate vs. multidimensional connectivity

We wanted to test whether our new TL-MDPC approach indeed captures more information than a comparable univariate approach for realistic choices of number of trials, number of vertices and signal-to-noise ratios (SNR). Thus, we compared TL-MDPC with a univariate approach in which summarised activation values (e.g., averages across vertices per ROI) are regressed against each other for pairs of ROIs at different latencies. This is similar to the study of Anzellotti et al (2017a) for fMRI data in combination with prior data reduction via PCA. The authors argued that a fair comparison between a univariate and multidimensional approach should be based on explained variance on the same data. However, since the univariate approach collapses the multi-voxel data into a single time course, it has less data to explain than the multidimensional approach that attempts to explain the variance across all voxels. Therefore, the authors projected the data from the univariate approach back into the original multidimensional space before computing the explained variance. Here, we take a different approach. We argue that because we attempt to compare methods in the way they would be applied to real data, i.e., to find whether there is evidence for a connection and not to predict the particular activation value in each vertex, we do not have to back-project the univariate data. This may result in an advantage of one method over the other in specific situations – but this is what the methods are designed for. For example, we expect the univariate method to perform better when patterns are indeed purely related in a univariate manner. This provides a more conservative test of our TL-MDPC method, highlighting the particular situations where it provides an advantage for detecting relationships between regions and not simply for predicting individual vertex responses. The critical tests are therefore, whether the univariate approach can also detect multidimensional relationships, and whether the multidimensional approach is sensitive to univariate effects.

### 2.2 Simulations: Performance of the univariate and multidimensional methods across different types of relationships between activity patterns

In this section, we compare the univariate and multidimensional approaches on simulated data for three different scenarios: no relationship between patterns, a univariate linear relationship and a multidimensional linear relationship. From a mathematical point of view, the estimation of pattern transformations does not depend on the exact time lag between the patterns in ROI X and Y, but only on the structure and relationships between patterns in these matrices. Thus, we can test properties of our methods in simulations without explicitly simulating time courses. Instead, we simulated scenarios with different properties for *X* and *Y*. Therefore, we will use the terms MDPC and UVC for the multidimensional and univariate approaches, respectively. Scenario 1 is illustrated in Figure 2a, which shows two random patterns that are independent of each other and thus have no connectivity, meaning that they cannot be mapped to each other by any function: *Y*≠ *f*(*X*). Figure 2b shows UV effects in scenario 2, where all vertices of ROI *X* show the same time course *x*of size *n*_*t*_ × 1 plus noise: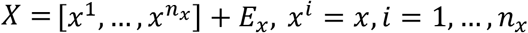. Vertices in ROI *Y*have the same time course *x*multiplied by a constant plus noise: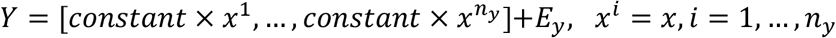. Scenario 3 in Figure 2c illustrates MD effects, with vertices in ROI X having no correlation with each other but being transformed to ROI Y through a matrix *T*, so that *Y*= *XT* + *E*, where *T* is of size *n*_*x*_× *n*_*y*_, and *E* is a zero-mean Gaussian matrix. For all cases, as in section 2.1.2, we predicted *Y*from *X* and vice versa, and took the EV average across the two directions as the connectivity metric.

**Figure 2.**
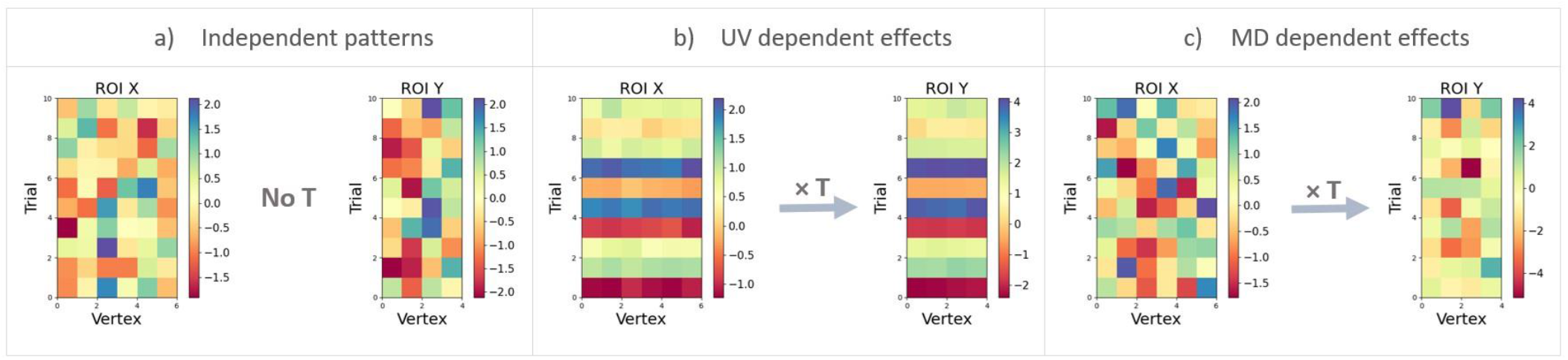
Illustration of different connectivity scenarios used in our simulations. The matrices illustrate activity patterns in regions *X* and *Y*, with rows representing different trials and columns representing vertices. a) *X* and *Y*contain random patterns. Thus, there is no reliable transformation between the patterns and consequently no connectivity. b) Patterns in *X* and *Y*have a univariate relationship. The amplitudes of patterns in the two regions covary linearly across trials (plus added noise). c) Patterns in *X* and *Y*have a multidimensional relationship. The patterns in *Y*were obtained by applying a linear transformation matrix to the patterns in *X* (plus noise).

#### 2.2.1 Simulation Parameters

In order to evaluate MDPC with respect to experimental parameters of practical relevance, we varied signal-to-noise ratio (SNR), number of vertices, and number of trials in our simulations within realistic ranges. As described above, we reduced the number of vertices per ROI using a clustering-based selection approach. In our real EEG/MEG data (below), this resulted in 5-13 vertices depending on ROI and latency range. We therefore manipulated the number of vertices in the simulations, including either 5 or 15 vertices in each region. For each set of parameters of each simulation, we computed the average of 100 simulations.

#### 2.2.2 Simulation 1: Checking for spurious connectivity measurement between two independent activity patterns

We first compared performance between MDPC and UVC approaches for the case where there is no relationship between ROI activity patterns (as in Figure 2a), in order to compare the false positive rates of the two methods. We generated two random noise patterns using normal distributions (mean=0 and std=1), and varied the number of trials [30, 50, 100, 150, 300] and vertices [5,15].

#### 2.2.3 Simulation 2: Testing the methods’ ability to detect the univariate dependency between two patterns

We then compared MDPC and the UVC method for the case where a univariate relationship exists between activity in ROIs X and Y (as in Figure 2b). For this purpose, we first created activity patterns for region X by generating one time course **x** from a normal distribution (mean=0, and std=1) and replicating this vector *n*_*x*_times (across vertices). To create patterns in ROI Y, **x** was multiplied by a constant and replicated *n*_*y*_ times. Again, different levels of noise were added using normal distributions (mean=0, and different standard deviations (std=10^std_pow^, where std_pow ∈[-2, -1.5, -1, -0.5, 0, 0.5, 1, 1.5])). As before, we varied the number of trials [30, 50, 100] and vertices [5, 15].

#### 2.2.4 Simulation 3: Testing the ability of each method to detect multidimensional connectivity between two patterns

In order to compare methods for multidimensional effects, we generated both patterns *X* and transformation matrix *T* using a normal distribution (mean=0, and std=1). *Y* was then obtained via a linear transformation of X by multiplying *X* by *T* (as illustrated in Figure 2c). We added different levels of noise from a normal distribution (mean=0) with a varying standard deviation (std=10^std_pow^, where std_pow ∈[-2, -1.5, -1, -0.5, 0, 0.5, 1, 1.5, 2])). We also varied the number of trials [30, 50, 100], as well as vertices [5, 15].

### 2.3 Real EEG/MEG dataset: Comparing TL-MDPC to a univariate approach through application to EEG/MEG data

As the next step, we asked whether and how the differences we observed between the MDPC and UVC approaches in simulated data manifest themselves in real data. Here the MDPC approach was the same as for the simulations, but applied over varying latencies, constituting the full TL-MDPC approach. Thus, we applied TL-MDPC and the time-lagged UVC (TL-UVC) approach to an existing dataset (Farahibozorg, 2018; Rahimi et al., 2022), aimed at revealing the task modulation of semantic brain networks through contrasting a semantic decision requiring deep semantic processing with a lexical decision task only requiring visual word recognition. Specifically, we asked whether the TL-MDPC approach identified connections within the dynamic semantic brain network that are missed with univariate approaches that cannot utilise the information contained in the patterns of activity within each brain region.

#### 2.3.1 Participants

We used data from 18 healthy native English speakers (mean age 27.00±5.13, 12 female) with normal or corrected-to normal vision. The experiment was approved by the Cambridge Psychology Research Ethics Committee and volunteers were paid for their time and effort.

#### 2.3.2 Stimuli and procedure

We used 250 words and 250 pseudowords in our analysis. The EEG/MEG experiment consisted of four blocks presented in a random order. One of the four blocks comprised a lexical decision (LD) task and the other three blocks comprised a semantic decision (SD) task. In the LD task, participants were required to identify whether the presented stimulus was referring to a meaningful word or a pseudoword. In the SD task, they were required to identify catch items where the presented word was referring to a specific group of words, “non-citrus fruits”, “something edible with a distinctive odour” or “food that contains milk, flour or egg”. This requires deeper semantic processing than lexical decisions, and as such contrasting SD over LD is expected to identify greater involvement of the semantic network, as found in Rahimi et al. (2022). Only word stimuli (no pseudoword catch items) were included in the following EEG/MEG analyses. Each stimulus was presented for 150ms, with an average SOA of 2400ms.

#### 2.3.3 Data Acquisition and Pre-processing

We used the same dataset presented in Rahimi et al., (2022). MEG and EEG data were acquired simultaneously using a Neuromag Vectorview system (Elekta AB, Stockholm, Sweden) and MEG-compatible EEG cap (EasyCap GmbH, Herrsching, Germany) at the MRC Cognition and Brain Sciences Unit, University of Cambridge, UK. MEG was recorded using a 306-channel system that comprised 204 planar gradiometers and 102 magnetometers. EEG was acquired using a 70-electrode system with an extended 10-10% electrode layout. Data were acquired with a sampling rate of 1000Hz.

To filter noise generated by distant sources, we applied Maxwell-Filter software to the raw MEG data (Taulu and Kajola, 2005). The preprocessing and source reconstruction were done in the MNE-Python software package (Gramfort et al., 2014, 2013). We visually inspected the raw data for each participant, and marked bad EEG channels for linear interpolation. We then used a finite-impulse-response (FIR) filter between 0.1 and 45 Hz. To remove artefacts (e.g. eye movement and heart signals), we applied the FastICA algorithm (Hyvarinen, 1999; Hyvärinen and Oja, 2000) and selected artefact components based on their temporal correlations with EOG signals. After ICA, data were divided into epochs from 300ms pre-stimulus to 600ms post-stimulus.

#### 2.3.4 Source Estimation

To reconstruct the source signals, we employed L2-Minimum Norm Estimation (MNE) (Hämäläinen and Ilmoniemi, 1994; Hauk, 2004). We then assembled inverse operators based on a 3-layer Boundary Element Model (BEM) of the head geometry obtained from structural MRI images. To do so, we assumed sources are perpendicular to the cortical surface (“fixed” orientation constraint). The noise covariance matrices were computed using baseline intervals of 300ms choosing the best based on cross-validated Gaussian likelihood from a list of methods from MNE Python (‘shrunk’, ‘diagonal_fixed’, ‘empirical’, ‘factor_analysis’) (Engemann and Gramfort, 2015). To regularise the inverse operator for evoked responses, we used MNE Python’s default SNR of 3.0. The participants’ source estimates were morphed to the standard Freesurfer brain (fsaverage).

#### 2.3.5 Regions of Interest

To examine the critical semantic network regions described in the Introduction, six regions of interest were selected including left and right anterior temporal lobes, left inferior frontal gyrus, left posterior temporal cortex, left angular gyrus, and left primary visual area (lATL, rATL, IFG, PTC, AG, and PVA) using the Human Connectome Project (HCP) parcellation (Glasser et al., 2016). See Rahimi et al. (2022) for more details.

#### 2.3.6 Applying TL-MDPC to the Real Brain Data

As described in 2.1.2, TL-MDPC can be computed for *X* predicting *Y* and for *Y*predicting *X*. Because we do not consider it meaningful to interpret the differences between these two cases, we averaged their results in the following analyses. Results were averaged across the two directions for every 25ms of data from 100ms pre-stimulus to 500ms post-stimulus. The connectivity score for each pair of ROIs at each latency, is presented in a TTM. Every row of a TTM shows dependencies between *Y* at a specific time point and *X* across the whole time period (across all columns), while every column indicates dependencies between *X* at a specific time point and *Y* over time (across all rows). Thus, we can explore statistical dependencies at different time lags.

Within a TTM we can distinguish three broad areas of interest:

1. *t*_*Y*_= *t*_X_: The diagonal shows simultaneous or zero-lag dependencies between *X* and *Y*.
2. *t*_*Y*_< *t*_X_: The lower triangle shows dependencies between current patterns of X and past patterns of Y, i.e. dependencies in which *X* is ahead of *Y*.
3. *t*_*Y*_> *t*_X_: The upper triangle shows dependencies in which *Y*is ahead of *X*.

Points 2) and 3) indicate that the upper and lower triangles capture different information. Consequently, TTMs are not necessarily symmetrical.

As in the simulations, the TTMs were masked to replace negative values with 0. Furthermore, to ensure connectivity estimation is not biased due to the different numbers of trials between our two tasks (Bastos and Schoffelen, 2016), the final TTMs were computed individually for the three SD blocks and the results averaged before comparison with LD, similar to our previous study (Rahimi et al., 2022). We then compared the TTMs of SD and LD using cluster-based permutation tests, implemented in MNE Python (Maris and Oostenveld, 2007), accounting for multiple observations (participants) across the different latencies of both ROI Y and X. To do so, t-values were computed and thresholded with a t-value equivalent to p-value < 0.05 for a given number of observations, and randomization was replicated 5000 times to determine the largest cluster size likely to be identified in data without true differences between the conditions. We applied two-tailed t-tests and the upper (SD>LD) and lower (LD>SD) 2.5% values in the resulting permutation distribution were considered to be significant. In order to remove small and possibly spurious clusters, we only reported clusters whose size was greater than 2% of the TTMs total size (in this case 0.02*(24*24) ≈ 12).

## 3 Results

### 3.1 Simulation results

#### 3.1.1 Simulation 1: Checking for spurious connectivity measurement between two independent activity patterns

We first assessed the case where there is no (univariate or multivariate) relationship between the activity patterns in two simulated ROIs (Figure 2a). In this scenario, both the MDPC approach and the UVC comparison approach should fail to identify a connection unless they are prone to false positive errors. Figure 3a shows the connectivity metric (explained variance, y-axis) for different number of trials (x-axis) and different numbers of vertices using the UVC approach. All values are close to zero, indicating that the UVC method does not produce false positive connections between two independent patterns. The same holds for the MDPC approach in Figure 3b. Thus, neither the MDPC nor the UVC approach are prone to yielding spurious connectivity for independent patterns.

**Figure 3.**
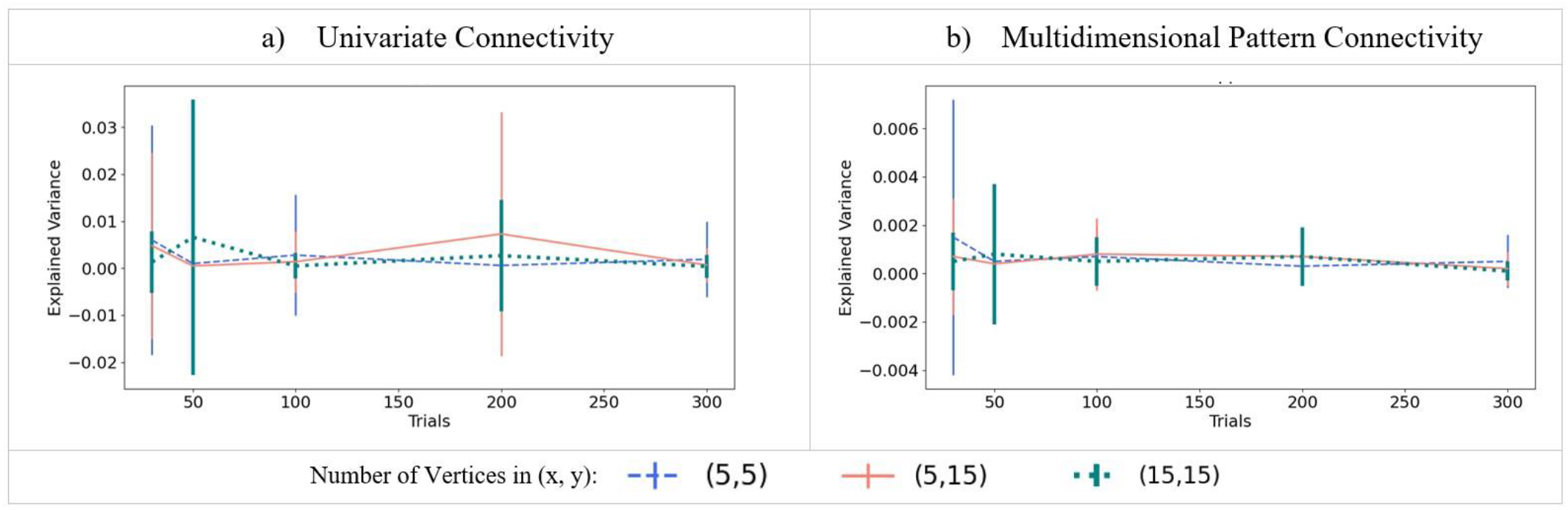
Estimating connectivity (explained variance) between two independent activity patterns, i.e., simulated regions without true connectivity, using MDPC and the UVC approach. a) Connectivity measured using the UVC method as a function of the different numbers of trials, for three different combinations of number of vertices in ROI X and ROI Y. b) Similar to a), but for the MDPC method. Both UVC and MDPC methods show connectivity scores close to zero, i.e., they do not produce false positives for independent patterns in the two regions. Error bars reflect standard deviations.

#### 3.1.2 Simulation 2: Testing the methods’ ability to detect univariate dependencies between two patterns

We next assessed how well the MDPC and UVC approaches could identify connectivity between activity patterns with a univariate dependency (as illustrated in Figure 2b). The left-hand panels in Figure 4 show how well the UVC method (red) and MPDC (blue) methods perform when the simulated patterns have a univariate relationship. The same pattern can be seen across panels a, c and e indicating that different numbers of vertices in X and Y regions do not have a strong effect on the methods’ performance in this range. In each, EV approaches 1 for SNRs greater than 10db, and is almost zero for SNRs below -25db. For intermediate SNRs both UVC and MDPC produce above-zero explained variance, with higher values for UVC compared to MDPC. Importantly, the MDPC is able to capture the univariate dependency between the patterns at medium to high SNRs. As expected, this is also possible with the UVC approach. Thus, both methods can identify connectivity between regions which display a univariate relationship. Both methods demonstrate improved detection of the relationship from 30 to 50 trials, but hardly any improvement from 50 to 100 trials.

**Figure 4.**
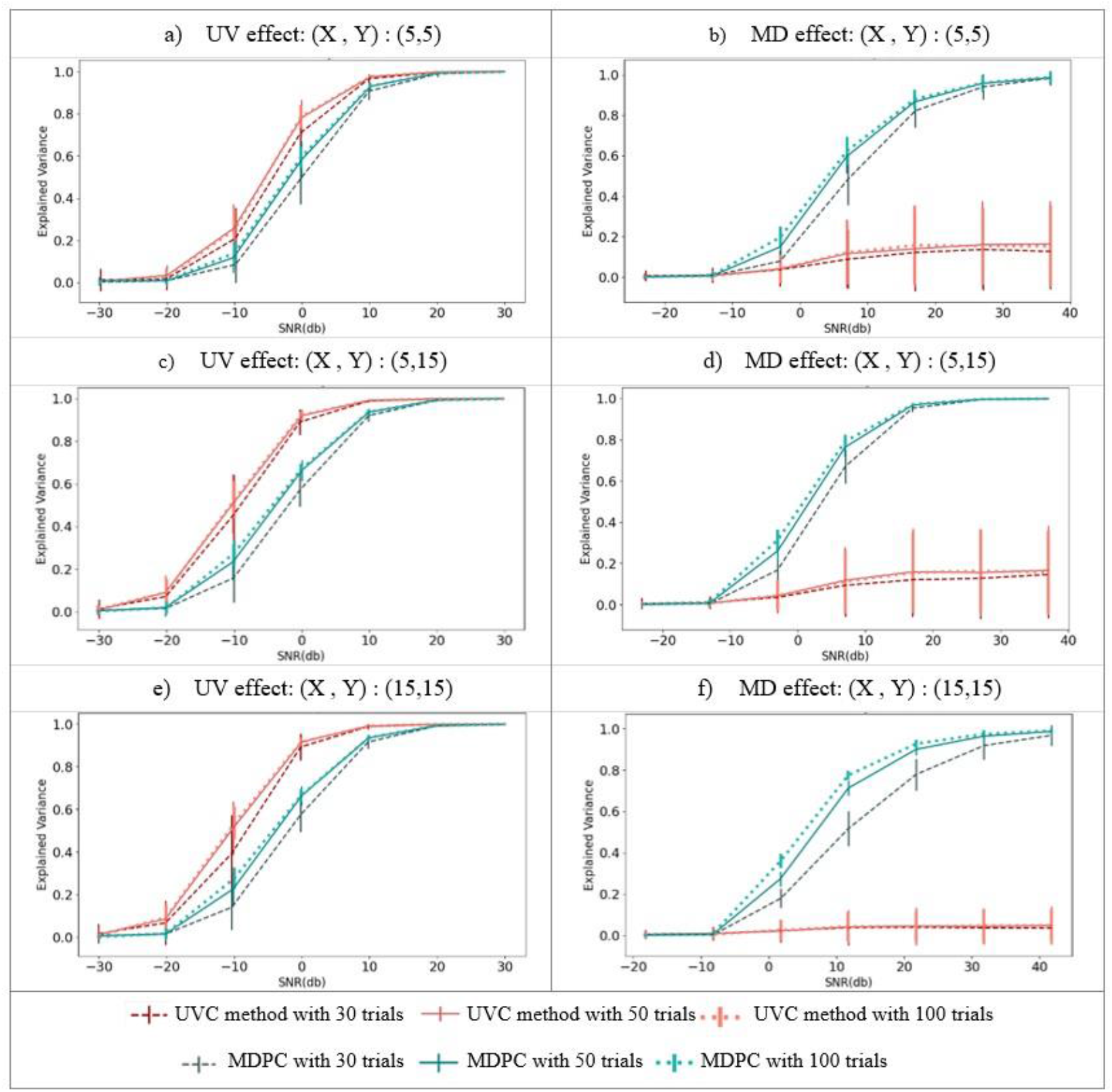
Comparing the MDPC and univariate connectivity (UVC) approaches on the detection of univariate (left panels) and multidimensional (right panels) relationships. All panels show explained variance (y-axis) across different SNRs (x-axis) for the MDPC (blue) and UVC (red) approaches with different numbers of trials (darker versus lighter shades of blue or red). a, c, e) show the connectivity scores for simulated activity patterns with a univariate dependency. In all cases, EV approaches 1 for SNRs greater than 10db, and is almost zero for SNRs below -25db. Both UVC and MDPC methods capture univariate relationships in medium-to-high SNRs, with greater EV for UVC. b, d, f) represent connectivity scores between patterns with a multidimensional dependency. For the MDPC method, EV approaches 1 for SNRs greater than 20db, and is almost zero for SNRs below -10db. For the UVC approach, the EV never exceeds a variance of 0.2, and is almost zero below -10db. Error bars reflect standard deviations.

#### 3.1.3 Simulation 3: Testing the ability of each method to detect multidimensional connectivity between two patterns

In a final set of simulations, we assessed how well the MDPC and UVC methods are able to detect a multidimensional relationship between two regions (illustrated in Figure 2c). The results for the MDPC (blue) and the UVC (red) approaches are displayed in the right-hand panels of Figure 4. Whilst connectivity scores for the MDPC increase gradually and eventually reach 1 at high SNRs (> 20db), the UVC approach fails to capture the multidimensional dependency and cannot explain more than 0.2 of data variance. Regardless of the number of vertices or trials, the MDPC method outperforms the UVC approach for SNRs > -5db (below which both methods perform poorly). The MDPC captures the multidimensional relationships between activity patterns, which are missed with the UVC approach. As with the UV relationships the MDPC performance increases with more trials, with larger improvement from 30 to 50 than from 50 to 100 trials.

Overall, our simulations demonstrated that both the MDPC and UVC approaches avoid false positives, whilst identifying univariate pattern dependencies. However, only the MDPC can capture multidimensional pattern dependencies between regions. Thus, the MDPC may identify additional connections between brain areas that would typically be overlooked with standard UV approaches.

### 3.2 Real EEG/MEG dataset: Comparing TL-MDPC to a univariate approach through application to EEG/MEG

In this section, we applied the same methods to a real EEG/MEG dataset to test whether this ability to identify additional, multidimensional relationships enables the TL-MDPC approach to uncover additional dynamic connections within the brain. We identify differences in connectivity within the semantic network when greater semantic processing is required, by contrasting a semantic decision task with a lexical decision task. A detailed analysis of the task modulation of evoked responses and univariate functional connectivity (measured as coherence) in this dataset can be found in our previous publication (Rahimi et al., 2022). Here, we will test whether TL-MDPC identifies additional connectivity when applied to this dataset, highlighting the advantages of this novel connectivity method over univariate approaches and extending our previous assessment of dynamic connectivity within the semantic network. Figure 5 illustrates the application of TL-MDPC and the time-lagged UVC approach to EEG/MEG data for two ROIs, namely lATL and IFG, key regions involved in semantic representation and control, respectively. Example TTMs are presented for the univariate analysis (panel a) and TL-MDPC (panel b), for the semantic decision (SD) and lexical decision (LD) tasks, as well as their statistical comparison (with cluster-based permutation tests). Greater semantic demands result in greater connectivity between lATL and IFG, a pattern which can be seen in both the TL-UVC and TL-MDPC analyses. However, this difference is only significant at a limited number of early time points in the TL-UVC case with short latencies. In contrast, the TL-MDPC approach identifies task-dependent semantic connectivity between lATL and IFG throughout the trial, and can detect relationships over longer latencies.

**Figure 5.**
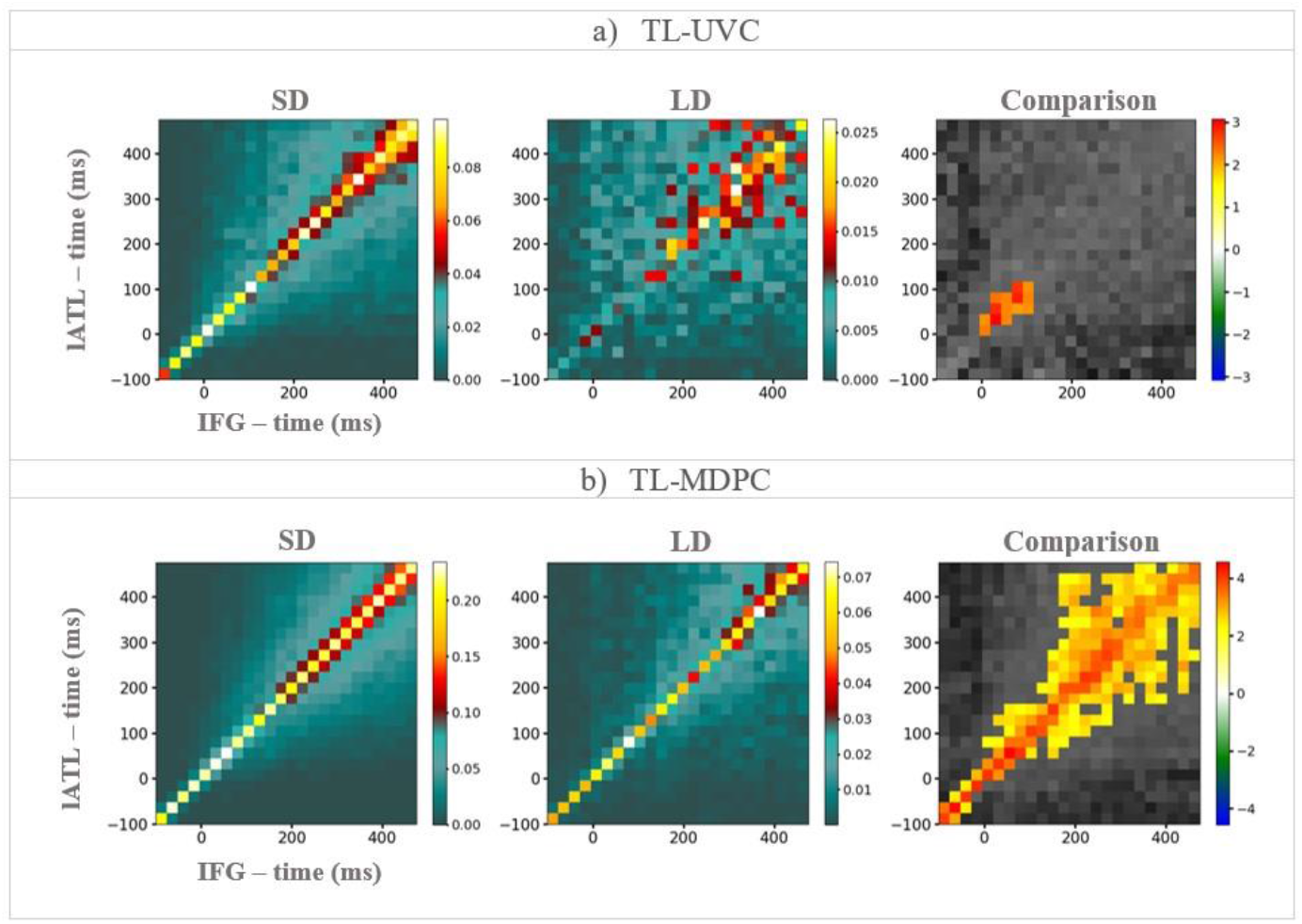
An example of TTMs showing the connectivity between the left ATL (y-axis) and IFG (x-axis), for the semantic decision (SD) task (the left column), the lexical decision (LD) task (the middle column), and their comparison (the right column) for which cluster size was thresholded at 12 cells. a) TTMs obtained using the time-lagged univariate connectivity (TL-UVC) approach demonstrate some modulation between the two tasks at a limited number of time points early in processing. b) TTMs obtained using the TL-MDPC method. Here, there is a wider pattern of significant task differences across the trial with prediction possible across longer lags. The upper diagonal indicates statistical dependencies where the lATL is ahead in time, and the lower part shows dependencies where the IFG is ahead in time, capturing different information. Colorbars show connectivity scores for the first two columns and t-values for the third column. Note, the matrices have idiosyncratic scales for display purposes, and the connectivity values are greater for the TL-MDPC method than the TL-UVC approach.

For both methods, the largest connectivity values occur along the matrix diagonal, i.e., at zero lag. For the TL-MDPC approach, differences are present in this example even in the baseline interval. This could be due to leakage or true baseline differences between our tasks, a possibility we discuss further below. When applying the TL-MDPC approach to both tasks, off-diagonal connectivity values start to fan out after stimulus presentation at 0ms, with the area of larger values broadening over time. The same pattern of increasingly broad off-diagonal connectivity is identified by the statistical analysis of the task differences presented in the right-most panel of Figure 5b. The pattern around the diagonal is approximately symmetric, suggesting that there is a statistical relationship between the patterns in the two regions, which varies depending on the pair of latencies involved, but not their order. For example, if there is a relationship between X at 200ms and Y at 300ms, then there is typically also a relationship between X at 300ms and Y at 200ms. TL-MDPC is a non-directional functional connectivity method and cannot be used to infer the directionality of the corresponding connections. How the temporal information identified can be used to characterize connectivity in more detail will be an important question for future research.

### 3.2.2 Capturing the connectivity within the semantic network across time with TL-MDPC

We next apply TL-MDPC to examine the connectivity between our full set of semantic ROIs, and compare these results to the TL-UVC approach. To summarise the TTM comparisons for all pairs of ROIs, we arranged the statistical results in a larger matrix, which we call an ‘inter-regional connectivity matrix’ (ICM), displayed in Figure 6. The upper diagonal (grey area) represents the transposed TL-UVC TTMs, and the lower diagonal (green area) shows TTMs based on the TL-MDPC results. Figure in the Supplementary Materials, provides the TTMs for each individual task for both TL-MDPC and univariate approaches. The most striking result is that TL-MDPC revealed task modulation of many more connections than the TL-UVC approach. For both approaches, all significant modulations show greater connectivity scores with a greater demand for semantic cognition. For the TL-UVC method, four connections were modulated by task demands: lATL-IFG, lATL-PTC, rATL-AG, and PTC-PVA with larger connectivity scores for SD.

**Figure 6.**
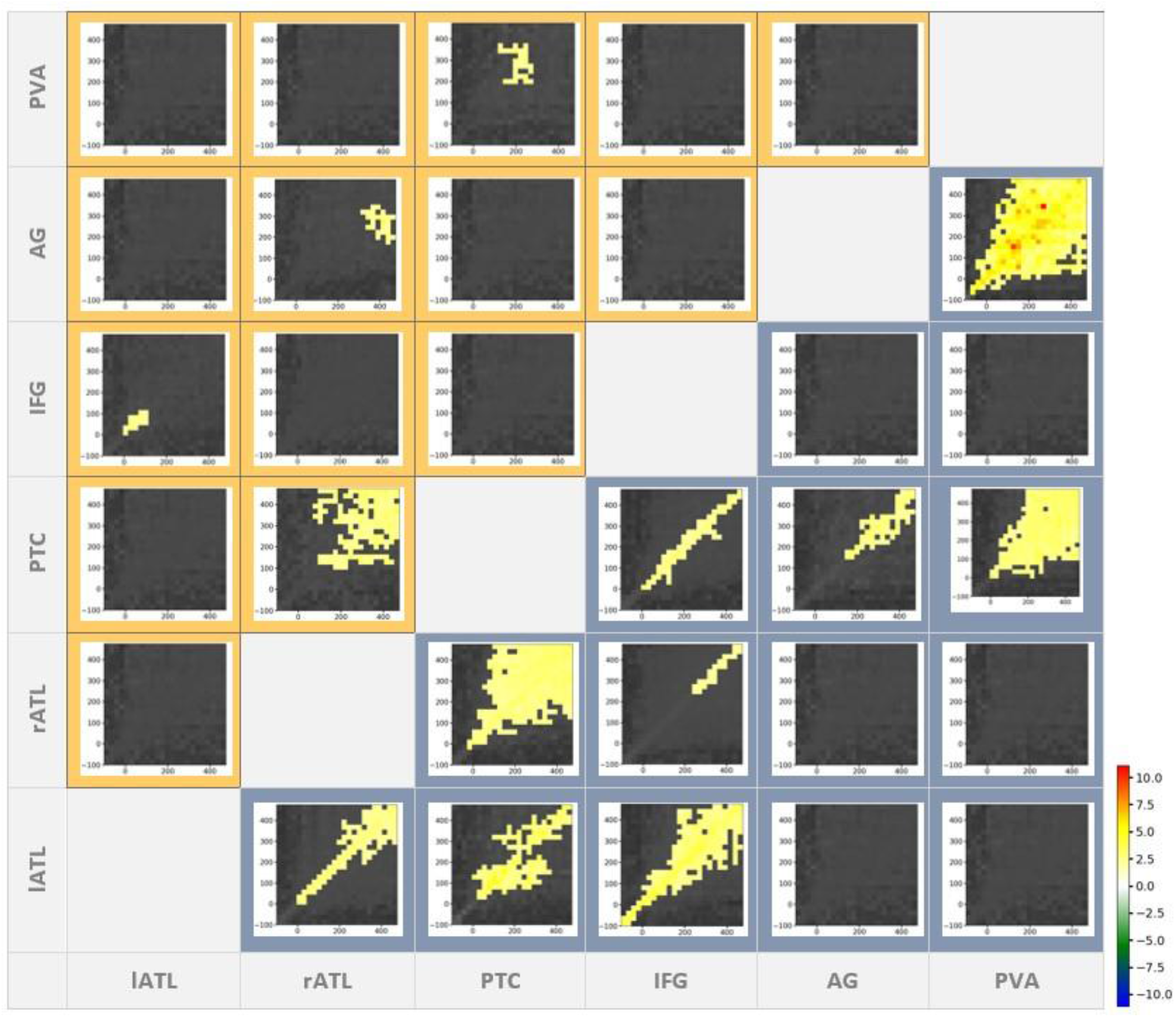
The Inter-regional Connectivity Matrix (ICM) – the upper triangle (yellow shaded area) shows temporal transformation matrices (TTMs) created using the TL-UVC approach and the lower diagonal (blue shaded area) TTMs constructed using the TL-MDPC approach. All TTMs shown are formed from the t-tests comparing the more over the less semantically demanding task. Cluster-based permutation tests were used for statistical comparisons. The alpha-level for both vertex-wise and cluster-wise t-tests was 0.05, and cluster size was thresholded at 12. The TL-UVC approach produced significant task modulation for four connections including lATL-IFG, lATL-PTC, rATL-AG, and PTC-PVA, with greater connectivity for SD. The TL-MDPC approach revealed a number of additional connections: lATL-rATL, lATL-PTC, lATL- IFG, rATL-PTC, rATL-IFG, PTC-IFG, PTC-AG, PTC-PVA, and AG-PVA, all with greater connectivity scores for SD than LD. The colour bar shows the scale of the t-values.

The TL-MDPC revealed modulations for nine ROI pairs, including: lATL-rATL, lATL-PTC, lATL-IFG, rATL-PTC, rATL-IFG, PTC-IFG, PTC-AG, PTC-PVA, and AG-PVA. Additionally, these modulations were significant across a larger time window. Our simulation results indicate that this increased sensitivity of TL-MDPC compared to TL-UVC approaches is unlikely to be due to false positives. Instead, the ability of the TL-MDPC approach to utilise the multidimensional activity pattern across a brain region may allow identification of connections that are missed with univariate connectivity approaches, as demonstrated with the simulated data. The current analyses highlight that this increased sensitivity to additional, multidimensional connections may extend to real EEG/MEG data, detecting plausible connectivity changes across known task networks. It is also possible that MDPC is more prone to leakage than UV, which we consider in the Discussion. It is noteworthy that although the ROIs included have all been linked to semantics, the connections revealed by TL-MDPC do not appear to be arbitrary. Instead, the core regions consistently implicated in multimodal semantic cognition, left and right ATL, IFG and PTC, show strong interconnectivity. In contrast, the PVA which is a visual region required to interact with many networks and not merely the semantic network, and the AG, which may have a role in some aspect of semantic or episodic memory, are not well connected to this core semantic network, but do connect to each other and the PTC. TL-MDPC appears to identify meaningful dissociations within the regions assessed. These connectivity differences are present at early time points and are prolonged across the trial, confirming our previous findings of semantic task modulations on evoked responses and functional connectivity in early processing stages (Rahimi et al., 2022).

As in the example in Figure 5, most TTMs show connections centred around the diagonal. Interestingly, some TTMs show patterns of significant effects diverging from the diagonal at early latencies, e.g., connections involving the visual cortex (AG-PVA and PTC-PVA), while others diverge later (e.g., PTC-rATL), and yet others stay around the diagonal (e.g., connections involving IFG). In some ROIs, significant zero-lag effects may be seen on the diagonal even in the baseline period. We must note that the significance of these effects was probed using cluster-based permutation tests, which means that determining their precise temporal extent (onsets and offsets) is not straightforward as the identification of an effect depends on the significance of contiguous cells of the matrix (Sassenhagen and Draschkow, 2019). This is a general issue, since the onset of effects in noisy data depends on both signal and noise (e.g. Hauk et al., 2012). However, our results suggest that there are task differences in the baseline, at least at an uncorrected significance level. It is possible that TL-MDPC is more prone to false positives than TL-UVC. However, our simulation results suggest that this is not the case, at least not for random patterns. Instead this may reflect real differences between the rest periods of our two tasks, which when presented in a blocked design, may indeed produce different baseline connectivity prior to stimulus onset as participants anticipate the upcoming stimulus.

We provide a summary of our results, also including findings from our previous study using coherence analysis for comparison, in Figure 7. Figure 7a and b show the network revealed in an early and late time window, respectively, using 1) coherence in our previous study (Rahimi et al., 2022), 2) the TL-UVC method tested as a comparison here, and 3) TL-MDPC. Both the current univariate analyses and the standard coherence metric identify very limited task-related connectivity changes in the semantic network, despite the large task-differences in evoked responses throughout the semantic network and the large change in the necessity of semantic cognition for the two tasks. In contrast, TL-MDPC demonstrates strong connectivity between regions across the semantic network. The majority of the connectivity changes highlighted with TL-MDPC, are present within the early time window (with the exception of rATL-IFG and AG-PTC). The identification of rich connectivity throughout the core semantic network, including key regions for semantic representation (left and right ATL) and semantic control (IFG and PTC) persists from the early to the late latency window. In contrast, the connectivity of the visual PVA region, and the AG whose role in semantic cognition is debated, show relatively sparse connectivity and do not connect to the core semantic network, with the exception of the PTC. This is true across time points with the AG only connecting to PVA at the early time point, and the additional connection to PTC apparent only at the later time point.

**Figure 7.**
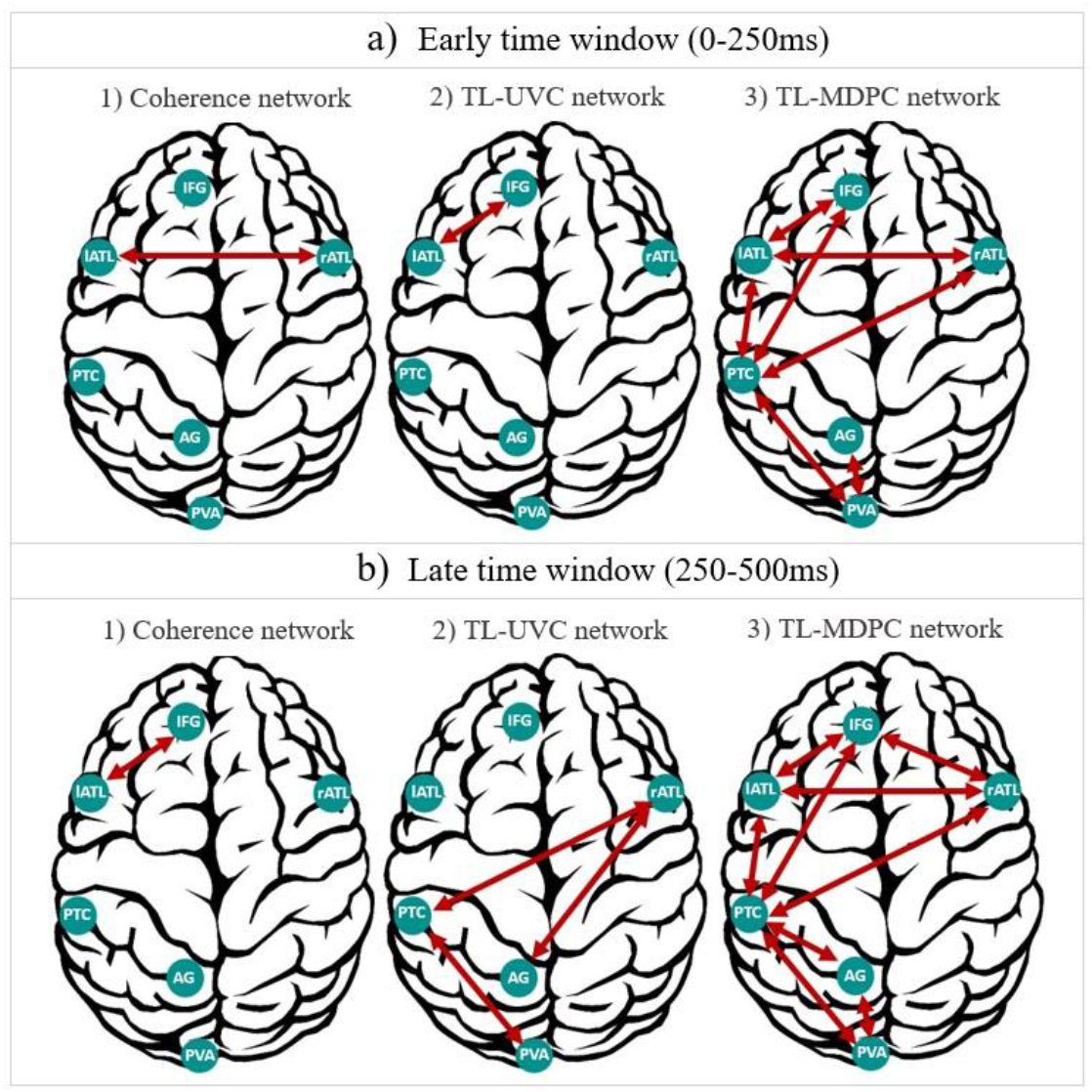
Summary of semantic task modulations of brain connectivity revealed by three different approaches. a) Networks revealed in an early time window (0-250ms), using 1) coherence (Rahimi et al., 2022), 2) TL-UVC method, and 3) TL-MDPC. b) Same as a), but for a later time window (250-500ms). In general, TL-MDPC captures many more connections compared to the other two methods. Importantly, for both time windows, TL-MDPC highlights rich connectivity between core semantic representation (lATL and rATL) and semantic control (IFG and PTC) areas, while AG connectivity is comparatively sparse, connecting to the visual sensory region but little of the core semantic network.

## 4 Discussion

In this study, we introduced time-lagged multidimensional pattern connectivity (TL-MDPC) to investigate the multidimensional relationships between patterns of event-related brain activation in space and time. TL-MDPC makes use of the full vertex-to-vertex transformations between patterns in different brain regions and is well-suited for EEG and MEG applications, exploiting their high temporal resolution to describe the bivariate relationships between patterns in pairs of ROIs across different pairs of latencies. We showed in simulations that neither a univariate (UVC) nor a multidimensional pattern connectivity approach (MDPC) are prone to false positives in the case of random and independent patterns in different regions. Not surprisingly, for simulated patterns that had a univariate (but no multidimensional) relationship between regions the UVC method outperformed the MDPC. In this case, the simulated scenario exactly matched the assumptions underlying the UVC method. Nevertheless, MDPC was able to capture univariate relationships at medium-to-high SNRs. In contrast, for simulated multidimensional pattern relationships the UVC method performed poorly even at high SNRs, while the MDPC performed well. Thus, TL-MDPC is sensitive to both univariate and multidimensional pattern relationships and may provide a more complete picture of brain connectivity than the univariate approach.

This pattern was confirmed in our analysis of real EEG/MEG data. TL-MDPC was able to identify task-dependent connectivity changes across the semantic network that the univariate approach and spectral coherence failed to detect, likely due to the presence of multidimensional connections. Using TL-MDPC, we observed rich connectivity between core semantic regions, including the left and right ATL hubs for semantic representation and the IFG and PTC regions crucial for semantic control. These changes were prolonged throughout the epoch and started early in processing. Not all regions assessed demonstrated such broad changes in connectivity throughout the semantic network, with PVA and AG demonstrating more limited connectivity changes. We conclude that TL-MDPC provides a richer description of the semantic brain network across time than univariate methods.

The description of the semantic network connectivity provided by TL-MDPC is highly informative with clear implications for the roles of the regions assessed and how they interact. The rich task-related connectivity between core semantic regions aligns with the central tenets of the controlled semantic cognition framework, supporting both the importance of the bilateral ATLs as hubs for multimodal semantic representation and the need for both control and representation regions (Lambon Ralph et al., 2016). Rahimi et al. (2022) delineated changes in evoked responses with greater semantic demand across this network, with strong differences in the ATLs. Both coherence and TL-MDPC identified stronger connectivity between left and right ATLs for the semantic compared to the lexical decision task. This corroborates the idea that right ATL is critical for semantic cognition and contributes more in semantically more demanding tasks (Rahimi et al., 2022; Stefaniak et al., 2022). Previous studies have showed that greater semantic demand leads to larger evoked responses in the IFG, primarily at later time points. This delay in engaging control areas, relative to representation regions was considered indicative that some initial representation may be performed without control, for instance, accessing some conceptual information before assessing how this information informs the task judgement. However, assessing task-related changes in connectivity with TL-MDPC identified early and persistent interactions between posterior temporal and inferior frontal control regions and the representational hubs. This demonstrates the importance of control areas throughout processing, suggesting that identification of task-relevant aspects of a concept is an iterative process requiring continual interaction between concepts and task context information, consistent with current models of controlled semantic cognition (Jackson et al., 2021). Gaining this additional understanding of how control and representation regions work together required the ability to track multidimensional relationships across time, demonstrating the utility of TL-MDPC of EEG/MEG data.

While rich connectivity changes were identified between the core semantic regions (lATL, rATL, IFG, PTC), the connectivity of the AG and PVA were relatively sparse. This finding may be expected for PVA, which is not responsible for multimodal semantic cognition but instead provides visual input to the semantic network, as well as to other networks responsible for other aspects of cognition. However, the relatively sparse connectivity of the AG with the core semantic network (in the context of early increases in evoked responses) may be more revealing. There are multiple possible explanations of this pattern of results: 1) AG may have some semantic-related role that is not part of the core semantic network, e.g., combinatorial semantics (Graves et al., 2010; Matchin et al., 2019; Price et al., 2016, 2015), 2) it could be performing a distinct task with the semantic stimuli, such as episodic encoding or directing attentional processes (Cabeza, 2008; Cabeza et al., 2012; Chambers et al., 2004; Humphreys et al., 2021; Humphreys and Lambon Ralph, 2015; Shimamura, 2011; Vilberg and Rugg, 2008; Wagner et al., 2005), or 3) task-related changes in the AG could simply reflect leakage from nearby visual areas which have a similar early time course resulting in connections with these nearby ROIs only. Overall, our findings provide little support for a core semantic role for the AG, but are compatible with a non-semantic role of the AG for example related to control and episodic memory (Farahibozorg et al., 2022; Humphreys et al., 2021; Noonan et al., 2013).

Intriguingly, both TL-MDPC and the univariate approach identified task modulations in connectivity prior to stimulus presentation. These effects had short lag times and could therefore reflect leakage, yet this would not explain why these effects are modulated by the task. Instead, as our tasks were administered in a blocked design, it is possible that task affected baseline activity, e.g. anticipatory processes or alpha rhythm. However, it is important to note that the extent of significant effects cannot be accurately inferred from cluster-based permutation testing (Sassenhagen and Draschkow, 2019), and onsets and offsets of effects are notoriously difficult to detect (Hauk et al., 2012). Our novel method offered the possibility to investigate the spatiotemporal dynamics in semantic brain networks in more detail than previous assessments utilizing fMRI data (Chiou et al., 2018; Jackson et al., 2016; Jung and Lambon Ralph, 2016) and univariate analyses (Rahimi et al., 2022; Sormaz et al., 2017). This demonstrates the clear utility of TL-MDPC to investigate the interactions within other cortical networks. One important methodological consideration for future applications of TL-MDPC is the sensitivity of the approach to leakage. In our previous study we provided a leakage matrix that describes the leakage among our ROIs based on point-spread and cross-talk functions (Rahimi et al., 2022). This analysis revealed that leakage among those ROIs was low to moderate. However, the leakage index used in that paper was based on the assumption of homogenous activation across vertices within ROIs and not complex patterns. Leakage from an active source occurs instantaneously, and is therefore often discussed in the context of zero-lag connectivity. However, since leakage occurs from each of a pair of ROIs, spurious connectivity can still be caused by leakage even at non-zero lags (Colclough et al., 2015; Farahibozorg et al., 2018; Palva et al., 2018). A more detailed analysis of leakage for multivariate and multidimensional scenarios should be performed in the future but is beyond the scope of the present study. Whilst increased leakage could lead to more connections as found with TL-MDPC, it is unlikely these results are due to leakage alone because of the following reasons. 1) The pattern of our results is meaningful in the context of current theories semantic brain networks. Our analysis distinguished between the core semantic network, comprising key semantic control and representation regions (lATL, rATL, IFG, and PTC) and the more restricted connectivity of posterior visual areas and the AG. 2) UV methods are not immune to leakage. The smoothness of patterns within ROIs should also result in some univariate leakage effects and affect the UVC method as well. Instead we observed meaningful and more widespread connectivity for TL-MDPC in line with our simulation results.

Further to introducing the TL-MDPC method, this study provides a novel approach for sub-sampling EEG/MEG data. EEG/MEG data contain redundant information due to their low spatial resolution, which can vary across ROIs. To alleviate this issue, we sub-sampled the most informative vertices within a brain region using a k-means clustering algorithm. This resulted in 5 to 13 vertices per region, suggesting a high degree of redundancy in the source estimates. It is well-established that the spatial resolution of source estimates is not only limited but also varies greatly across brain regions (e.g. is lower for deep rather than superficial locations) and depends on parameters such as the SNR (Hauk et al., 2019; Krishnaswamy et al., 2017; Samuelsson et al., 2021). In the future, our approach may become useful to quantify spatial resolution and the degree of redundancy across brain regions and for different parameter settings, especially for computationally demanding multivariate or multidimensional methods.

Our approach is a first step toward exploiting the full information contained in dynamic multidimensional data. Here, we presented a first application of the method proposed by Basti et al. (2019) and inspired by the work of Anzellotti et al. (2017a) for fMRI data to EEG/MEG data in source space. This initial method makes multidimensional connectivity available for EEG/MEG applications and opens multiple possibilities for extension in future work. For example, our approach could be extended to nonlinear pattern transformations, similar to the neural network approach by Anzellotti et al. (2017b). While linear methods can often produce reasonable approximations to nonlinear phenomena at least within certain parameter ranges, nonlinear methods are likely to complement linear approaches. However, there are many different ways to incorporate nonlinearities, and these methods may increase demands on numbers of trials and SNR. Therefore, this extension is non-trivial. Additionally, our proposed TL-MDPC approach is a bivariate non-directional functional connectivity method. It establishes statistical relationships between signals in two brain regions, but does not allow inferences about directionality or causality of effects. Our approach could be easily extended to autoregressive models that use a Granger-causality logic for multidimensional data (Granger, 1969; Hu et al., 2012). Finally, these methods could be applied to the time-frequency domain to reveal how time-frequency representations are transformed across brain regions. We hope that our study is a useful step towards future methods development and research that can “transform” spatiotemporal connectivity analyses.

## 5 Acknowledgements

This work was supported by intramural funding from the Medical Research Council UK (MC_UU_00005/18), a British Academy Postdoctoral Fellowship awarded to R.L.J. (No. pf170068), and Cambridge University international scholarships awarded to S.R. and S.R.F. For the purpose of open access, the author has applied a CC BY public copyright license to any Author Accepted Manuscript version arising from this submission. We also thank Professor Rik Henson for useful comments on a previous version of this manuscript.

## Data and code availability

Codes used for this study are available from the following public repository: https://github.com/setareh10/MDPC

## Credit author statement

**Setareh Rahimi:** Conceptualization, Methodology, Software, Formal analysis, Investigation, Visualization, Writing – original draft, Writing– review & editing, Project administration. **Rebecca Jackson:** Supervision, Conceptualization, Methodology, Investigation, Writing– review & editing, Project administration. **Seyedeh-RezvanFarahibozorg:** Data curation, Investigation, Editing. **Olaf Hauk:** Supervision, Conceptualization, Methodology, Software, Investigation, Writing– review & editing, Project administration, Resources

## Supplementary

**Figure S1.**
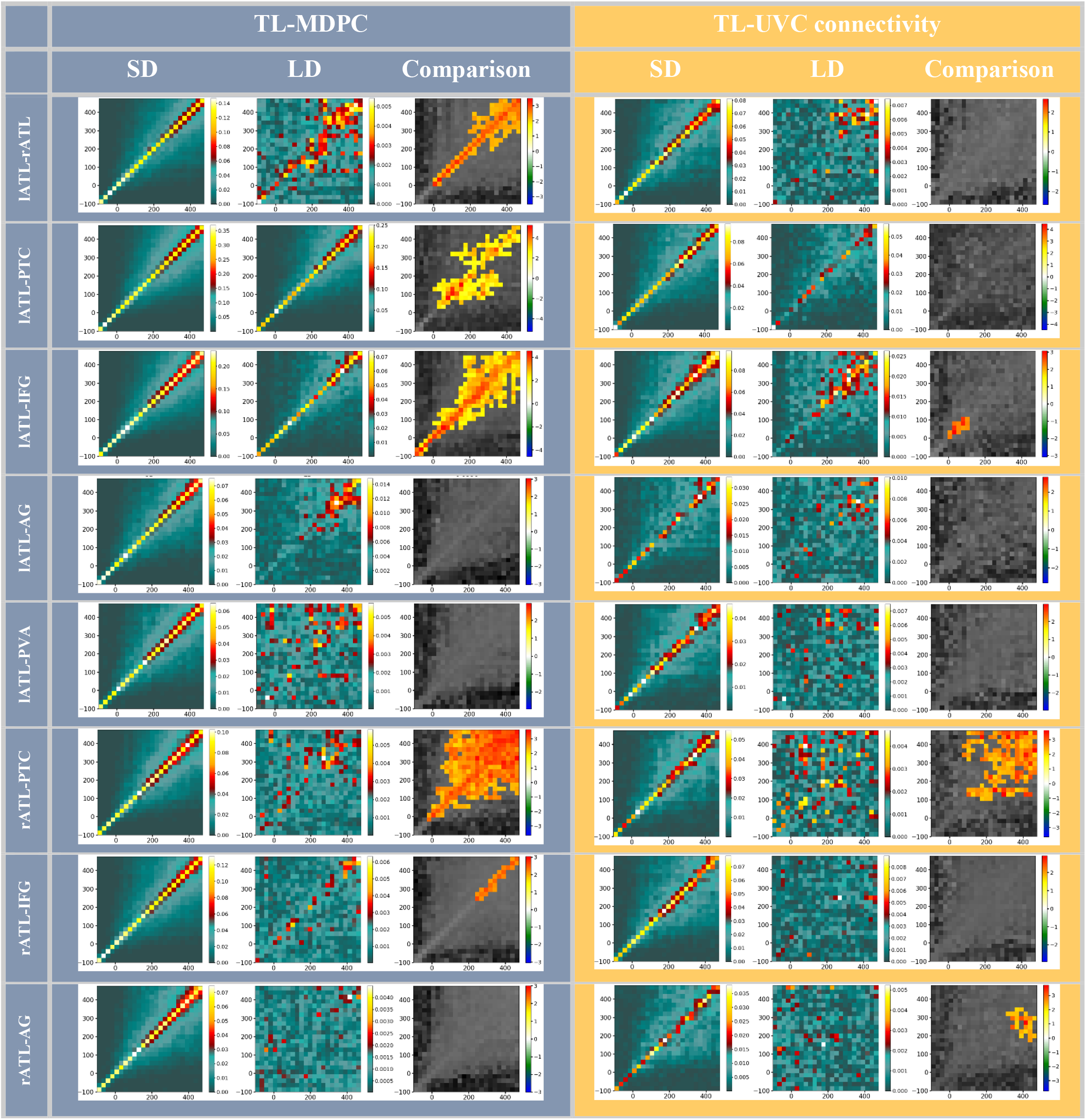
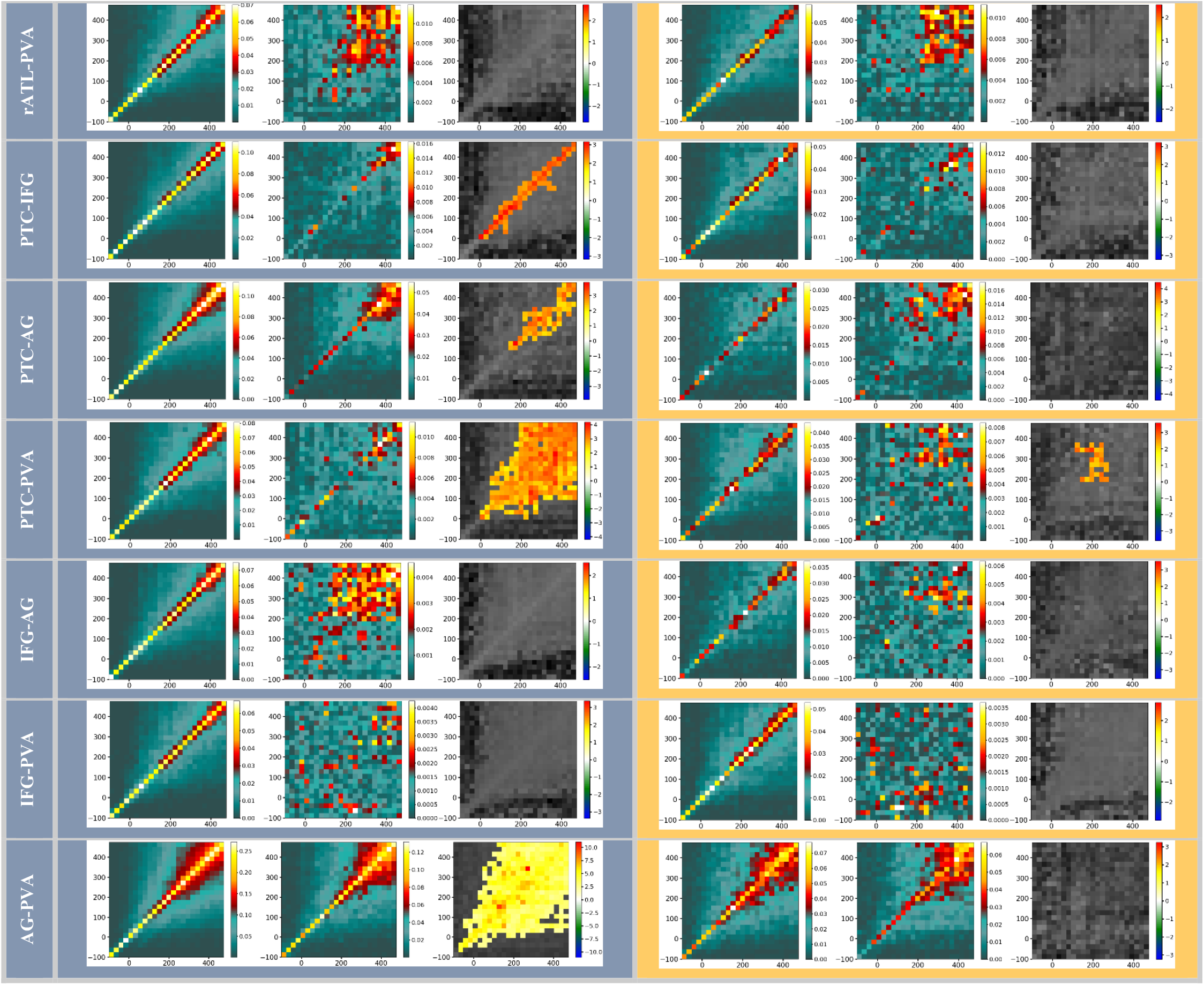
Representation of all TTMs for semantic decision (SD) and lexical decision (LD) tasks, and their comparison, using TL-MDPC (left column) and TL-UVC approach (right column). TTMs for SD and LD are averaged across participants, and comparison was performed using cluster-based permutation test with alpha-level=0.05. All significant contrasts show greater connectivity for SD using MD. The size of all clusters shown here is greater than 2% of TTMs size (24*24).

## Footnotes

1 https://www.scikit-yb.org/en/latest/api/cluster/elbow.html

